# Genomic surveillance for multidrug-resistant or hypervirulent *Klebsiella pneumoniae* among bloodstream isolates at a United States academic medical center

**DOI:** 10.1101/2020.12.04.410209

**Authors:** Travis J. Kochan, Sophia H. Nozick, Rachel L. Medernach, Bettina H. Cheung, Samuel W.M. Gatesy, Marine Lebrun-Corbin, Sumitra D. Mitra, Natalia Khalatyan, Fiorella Krapp, Chao Qi, Egon A. Ozer, Alan R. Hauser

## Abstract

**Background:** *Klebsiella pneumoniae (K. pneumoniae)* strains have been divided into two major categories: classical *K. pneumonia*, which are frequently multidrug-resistant and cause hospital-acquired infections in patients with impaired defenses, and hypervirulent *K. pneumoniae*, which cause severe disseminated community-acquired infections in immunologically normal hosts. Both types of infections may lead to bacteremia and are associated with significant morbidity and mortality. The relative burden of these two types of *K. pneumoniae* among bloodstream isolates within the United States is not well understood.

**Methods:** We examined consecutive *K. pneumoniae* isolates cultured from the blood of hospitalized patients at Northwestern Memorial Hospital (NMH) in Chicago, Illinois April 2015 – April 2017. Bloodstream isolates underwent whole genome sequencing, and clinically relevant data (sequence type [ST], capsule loci, virulence factors, and antimicrobial resistance genes) were inferred from the genome using the bioinformatic tools *Kleborate* and *Kaptive*. Patient demographic, comorbidity, and infection information, as well as phenotypic antimicrobial resistance of the isolate were extracted from the electronic medical record. Candidate hypervirulent isolates were tested in a murine model of pneumonia. Complete genome sequences of these candidate hypervirulent isolates were obtained and plasmid content was evaluated.

**Results:** *K. pneumoniae* bloodstream isolates (n=104) from NMH were highly diverse consisting of 75 distinct STs and 51 unique capsule loci. The majority of these isolates (n=58, 55.8%) were susceptible to all tested antibiotics except ampicillin, but 17 (16.3%) were multidrug-resistant. A total of 32 (30.8%) of these isolates were STs of known high-risk clones, including ST258 and ST45. In particular, 18 (17.3%) were resistant to ceftriaxone, 17 harbored extended-spectrum beta-lactamases, and 9 (8.7%) were resistant to meropenem. Four (3.8%) of the 104 isolates were hypervirulent *K. pneumoniae*, as evidenced by hypermucoviscous phenotypes, high levels of virulence in a murine model of pneumonia, and the presence of large plasmids similar to characterized hypervirulence plasmids. Of particular concern, several of these plasmids contained *tra* conjugation loci suggesting the potential for transmission. Two of these hypervirulent strains belonged to the well characterized ST23 lineage and one to the emerging ST66 lineage.

**Conclusions:** While NMH *K. pneumoniae* bloodstream infections are caused by highly diverse strains, the most prevalent STs were those of multidrug-resistant high-risk clones. A small number of hypervirulent *K. pneumoniae* were observed in patients with no recent travel history, suggesting that these isolates are undergoing community spread in the United States.

## Introduction

*Klebsiella pneumoniae* (*K. pneumoniae*) is a leading cause of nosocomial infections worldwide including pneumonia, bloodstream, and urinary tract infections (1). This bacterium causes severe secondary pneumonia in patients with respiratory virus infections and is a leading cause of secondary pneumonia in patients with COVID-19 (2). *K. pneumoniae* pathogenicity is due to a variety of virulence factors, including siderophores, fimbriae, and polysaccharide capsules of various types. The polysaccharide capsule allows this bacterium to evade host immune defenses, such as complement-mediated phagocytosis (3, 4).

Most *K. pneumoniae* are referred to as “classical” (cKP), which infect chronically ill patients residing in hospitals and long-term care facilities (5). Some cKP strains have become increasingly resistant to antibiotics, including carbapenems (6-11). A subgroup of cKP have been designated “high-risk clones,” lineages that are frequently multidrug-resistant (MDR) and extensively drug-resistant (XDR) and that have spread across continents to cause large numbers of infections (12). Some of the best characterized *K. pneumoniae* high-risk clones include ST11, ST14, ST15, ST17, ST147, ST258, ST307, and ST512. Within the United States, ST258 and ST307 are endemic high-risk clones that are carbapenem-resistant (CRE) and extended-spectrum-beta-lactamase-resistant (ESBL), respectively. Genes conferring antimicrobial resistance in these strains are often carried by conjugative plasmids that allow for dissemination among cKP strains. Thus, it is not surprising that the IDSA, WHO, and the CDC have each deemed MDR *K. pneumoniae* as a serious public health priority in need of new therapeutic development (13-15).

A second type of *K. pneumoniae*, referred to as hypervirulent *K. pneumoniae* (hvKP), was first identified in Taiwan in 1986 as a common cause of pyogenic liver abscesses (PLA) in young, otherwise healthy individuals living in the community (16-23). Other than diabetes mellitus, these patients lacked risk factors commonly associated with cKP infections. While these strains are now common in Asia, little is known about their prevalence in the United States. Case reports document the presence of hvKP infections in North America, but only a few surveillance studies have been performed (22, 24-26). hvKP strains frequently express highly mucoid capsules, leading to a “hypermucoviscous” (hmv) colony phenotype, and are considerably more virulent in mouse models than cKP strains (27). Their increased virulence is usually attributed to the presence of one of several large, non-conjugative “hypervirulence” plasmids (28-30). The non-conjugative nature of these plasmids and the limitations imposed by the hyper-mucoid capsule on genetic exchange may restrict the hypervirulence phenotype to strains of a few sequence types (e.g. ST23, ST86, ST66) and capsule loci (e.g. KL1, KL2)(18, 31, 32). Detection of hvKP is clinically important, as they may cause infections that require prolonged antibiotic therapy, that tend to relapse, and that frequently disseminate to remote sites (33).

Hypervirulence plasmids are defined by several pathogenic features. In general, these plasmids contain two distinct pathogenicity loci (PAL-1 and PAL-2) (Fig 1A). PAL-1 usually consists of a mucoid regulator gene (*rmpA2*), the aerobactin receptor gene (*iutA*), and aerobactin biosynthesis genes (*iucABCD*). PAL-2 usually consists of a distinct mucoid regulator operon (*rmpADC*), the salmochelin receptor gene (*iroN*), and salmochelin biosynthesis genes (*iroBD*) (Fig 1A). However, considerable diversity exists among individual hypervirulence plasmids (30, 34). For example, pK2044 and KP52.145pII, two of the most common and best characterized hypervirulence plasmids, differ in that KP52.145pII contains an incomplete PAL-1 that lacks *rmpA2* (Fig 1A, B, C). We previously identified plasmid pTK421_2 from a *K. pneumoniae* bloodstream isolate from Northwestern Memorial Hospital (NMH) in Chicago, Illinois (Fig 1D) (35). This plasmid also lacks *rmpA2* but contains conjugation genes (Fig 1D). The population diversity of hypervirulence plasmids remains poorly defined but is an area of active investigation.

**Figure 1.**
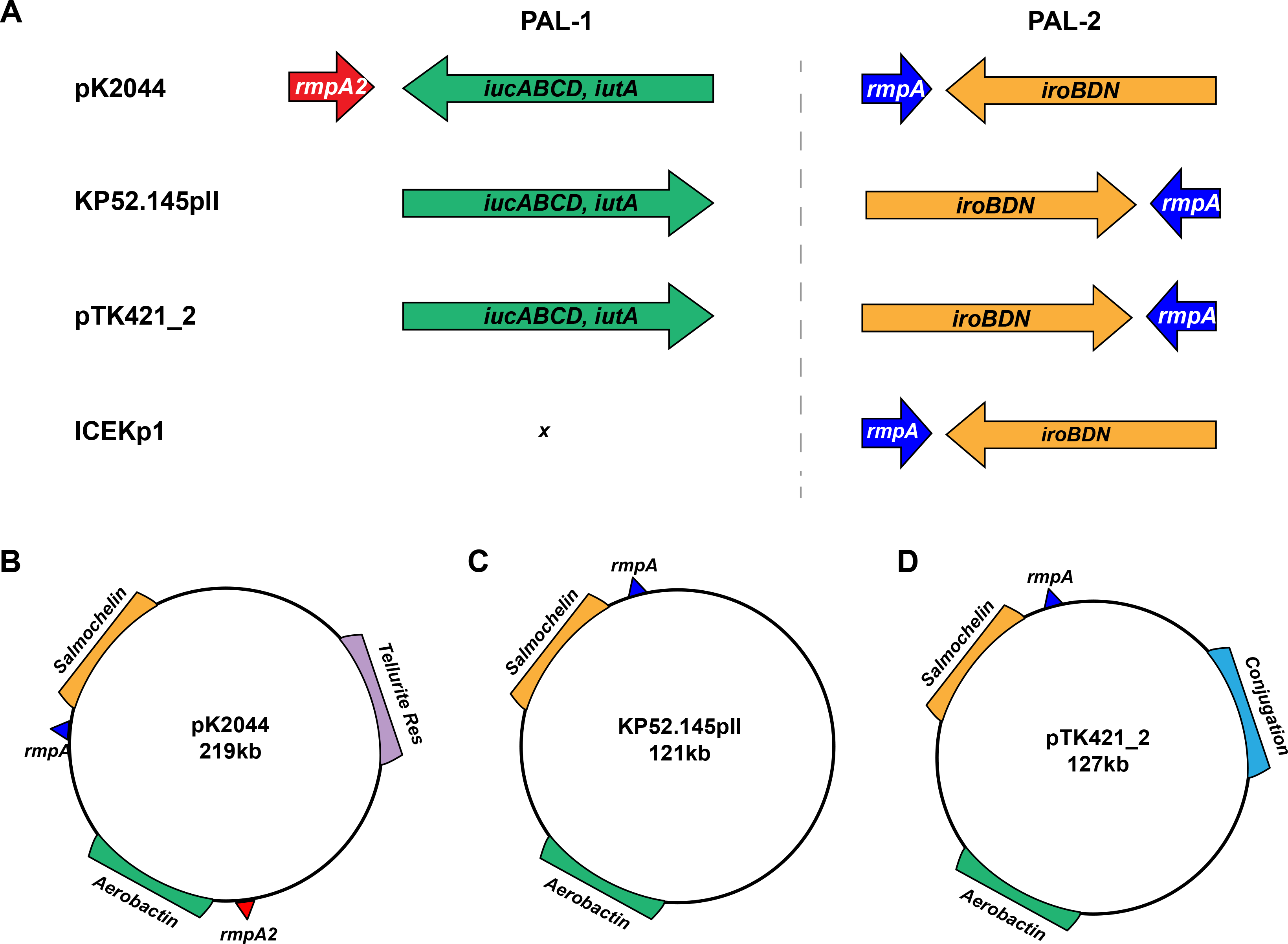
Virulence plasmids associated with hypervirulence in *Klebsiella pneumoniae*. PAL-1 and PAL-2 contents and organization are depicted for pK2044, KP52.145pII, pTK421_2, and ICEKp1 **(A)**. Genomic elements and plasmid architecture are depicted for pK2044 **(B)**, KP52.145pII **(C)** and pTK421 **(D)**. (Drawings not to scale.)

The hypervirulence plasmids contribute to the pathogenicity of hvKP strains in several ways. The mucoid regulators *rmpA* and *rmpA2* increase capsule production and hypermucoviscosity (29), and the siderophores are thought to provide access to host- sequestered iron reserves. However, the relative roles of the different accessory siderophores (aerobactin, salmochelin, and yersiniabactin) in hvKP virulence remain controversial (36, 37). A 2015 report suggested that aerobactin was the only siderophore required for the hypervirulence phenotype of the ST86-K2 capsule type strain hvKP1 in both mouse intraperitoneal and pneumonia models (37). In contrast, another report showed that only a triple siderophore mutant (yersiniabactin, aerobactin, and salmochelin) of the ST23-K1 strain NTUH-K2044 was attenuated in the mouse intraperitoneal model (36). In addition, strain ATCC43816 (also called KPPR1) is hypervirulent in a mouse model of pneumonia but is otherwise atypical in that it lacks a hypervirulence plasmid and PAL-1 but contains a chromosomal version of PAL-2 in an integrative and conjugative element (ICEKp1, Fig 1A) (4, 38). In KPPR1, yersiniabactin promoted disease in the lung and salmochelin promoted dissemination to tissues (39). Thus, it is clear that mucoid regulators and siderophores play roles in conferring the hypervirulent phenotype, but significant strain-to-strain variability may occur.

As mentioned, hvKP strains were first identified based upon the unusual disease manifestations they cause (e.g. PLA), but considerable effort has been devoted to developing a microbiological definition that could be used by clinical laboratories to identify these strains. There are three characteristics commonly associated with hvKP: 1) hypermucoid capsules, 2) presence of hvKP pathogenicity loci, and 3) high levels of virulence in mouse models of infection (5, 16, 18, 27, 31, 33, 40-45). Initially, hypermucoviscosity was used as a proxy for the hvKP phenotype. However, several reports have described hmv *K. pneumoniae* strains that lack pathogenicity loci usually associated with hvKP and are not hypervirulent in mice (46-48). As a result, genetic biomarkers have been used to define hvKP, but these definitions have varied from study to study. Recently, Russo and colleagues systematically examined the accuracy of using *rmpA*, *rmpA2*, *peg-344*, *iucA*, and *iroB*, as diagnostic biomarkers for hvKP (27, 49). In their cohort of 175 isolates, the presence of *iucA*, *iroB* or a hmv phenotype distinguished hvKP from cKP with an accuracy of 96%, 97%, and 90%, respectively.

We characterized 140 consecutive bloodstream isolates of *Klebsiella* from patients at NMH in Chicago, Illinois from 2015 – 2017. We sequenced and determined virulence gene content, antimicrobial resistant gene content, and hypermucoviscosity of these isolates. We identified numerous MDR cKP high-risk clones and several isolates that contained genomic, clinical, and virulence properties consistent with hvKP.

## Results

### Microbiological characteristics of *Klebsiella* bloodstream isolates

We collected 140 consecutive *Klebsiella spp.* isolates from 119 patients with positive blood cultures from 2015–2017 at NMH. The Clinical Microbiology Laboratory at NMH identified these isolates as *K. pneumoniae*. Note that the *Klebsiella pneumoniae* species complex has recently been subdivided into *Klebsiella variicola, Klebsiella quasipneumoniae subsp. quasipneumoniae*, *Klebsiella quasipneumoniae subsp. Similipneumoniae* (50). Thus, we performed Illumina whole genome sequencing of these presumptive *K. pneumoniae* bacteremia isolates to more definitively define their species. Among these isolates, 85% (n=119) were confirmed by whole-genome sequencing to be *K. pneumoniae* with the remaining isolates being *K. variicola* (n=16), *K. quasipneumoniae subsp. quasipneumoniae* (n=1), *K. quasipneumoniae subsp. similipneumoniae* (n=3), and *K. oxytoca* (n=1) (Fig S1). The 119 *K. pneumoniae* isolates were cultured from 101 patients and were highly diverse and consisted of 75 unique sequence types (STs) (Simpson’s diversity index = 0.98), 51 unique capsule loci (KL) (Simpson’s diversity index = 0.97), and 10 distinct O antigen serotypes (Simpson’s diversity index = 0.69). After removing isolates of the same strain cultured repeatedly from the same patient, we determined that 104 *K. pneumoniae* strains were obtained from 101 patients. Three patients had two separate *K. pneumoniae* bloodstream infections caused by distinct strains, which were included in this study. This collection includes several STs associated with carbapenem resistant high-risk clones such as ST258 (n=6), ST45 (n=6), ST29 (n=5), ST14 (n=3), ST17 (n=3), ST15 (n=3), ST147 (n=3). Two isolates belonged to the ESBL-containing high risk clone ST307 (Fig 2 & S2A). In addition, we identified three STs associated with hvKP: ST23 (n=2), ST66 (n=1), and ST380 (n=1) (Fig 2 & S2A). The most common KL-types were KL2 (n=7), KL107 (n=7), KL30 (n=6), KL25 (n=5), KL24 (n=4), KL28 (n=4), KL22 (n=4), and KL54 (n=4) (Fig S1B). The most common O antigen serotypes were O1 (n=38), O2 (n=27), and O3b (n=21) (Fig S2C). These findings indicate that bloodstream isolates at NMH are highly diverse with respect to their STs, capsule types, and O-antigen serotypes.

**Figure 2.**
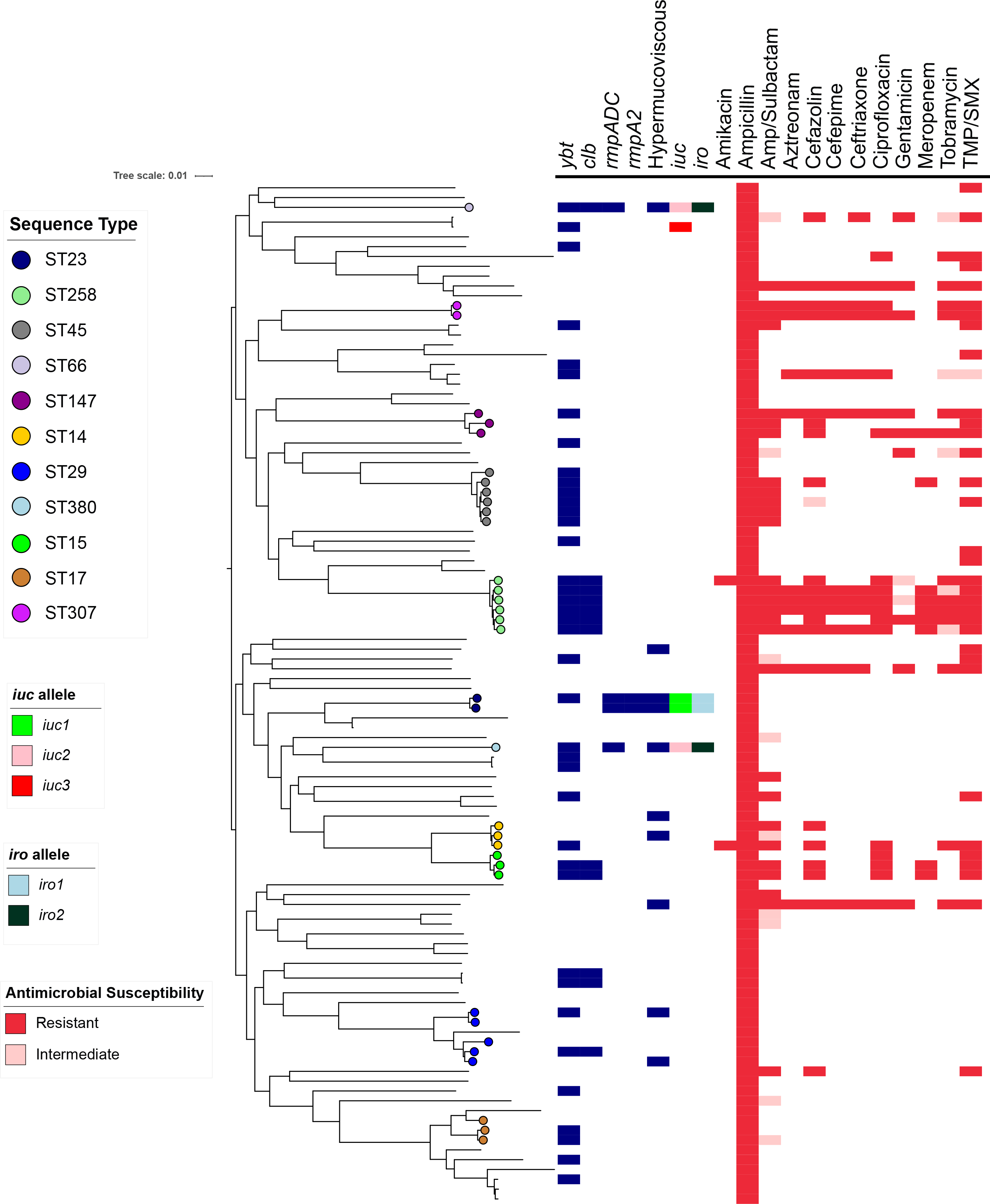
Core genome phylogenetic tree of 104 *Klebsiella pneumoniae* bloodstream isolates. Maximum likelihood phylogenetic tree generated from core genome SNP loci in 104 *K. pneumoniae* bloodstream isolates. Sequence types, the presence of virulence factors, and antibiotic resistances are indicated. *ybt* = yersiniabactin biosynthesis loci, *clb* = colibactin biosynthesis loci, *rmpADC* = mucoid regulator operon, *rmpA2* = regulator of mucoid phenotype 2, *iuc* = aerobactin biosynthesis genes, *iro* = salmochelin biosynthesis genes, Amp = ampicillin, TMP/SMX = trimethoprim-sulfamethoxazole. Strains were considered Hypermucoviscous if they were positive by string test. The complete genome of NTUH-K2044 is used as the reference genome.

### Antimicrobial Resistance Determinants

While many cKP are MDR, hvKP isolates tend to be susceptible to antibiotics. We therefore examined the antibiotic resistance profiles of the isolates in our collection. Since *K. pneumoniae* is inherently resistant to ampicillin, it was not surprising that 100% of the bloodstream isolates had MICs above the resistance threshold (Fig 2 & S3). Thirty-nine isolates (37.5%) were non-susceptible (29 resistant, 10 intermediate) to ampicillin/sulbactam, 17 (16.3%) were resistant to aztreonam, 23 (22.1%) were non-susceptible (21 resistant and one intermediate) to cefazolin, 18 isolates (17.3%) were resistant to ceftriaxone, 17 isolates (16.3%) were resistant to cefepime, and nine isolates (8.7%) were resistant to meropenem (Table S1, Fig 2 and S3). Among NMH 104 bloodstream isolates, 42% contained at least one acquired antimicrobial resistance gene (Table S1). Twelve isolates (11.5%) contained extended spectrum beta-lactamase genes and nine (8.7%) contained a carbapenamase gene (8 *bla_KPC-3_*, 1 *bla_NDM-1_*) (Fig S4). *bla_NDM-1_* was found in an ST147 isolate, while *bla_KPC-3_* carbapenemase genes were found in ST258 (n=5), ST15 (n=2), and ST45 (n=1) strains (Table S1). One ST258 isolate, KPN88, did not contain a carbapenamase gene and was sensitive to several beta-lactams (Table S1 and Fig 2). The most common ESBL genes were *bla*_CTX-M-15_ (n=7, 6.7%) and *bla*_SHV-12_ (n=4, 3.8%) (Table S1 and Fig S4). Strains carrying *bla*_CTX-M-15_ were highly drug resistant including all beta-lactams tested except meropenem (Fig S4A-F). Strains carrying *bla*_SHV-12_ also carried genes predicted to confer carbapenem resistance (Fig 2 & S4A-F). Isolates containing carbapenamases and/or ESBLs also carried genes predicted to confer resistance to aminoglycosides, trimethoprim, sulfonamides, tetracycline, fluoroquinolones, and phenicols suggesting these strains contain one or more antimicrobial resistance plasmids (Table S1). The correlations between phenotypic beta-lactam resistance and the presence of beta-lactam genes was very strong. All isolates containing a carbapenemase gene were resistant to ampicillin/sulbactam, aztreonam, cefazolin, cefepime, ceftriaxone, and meropenem (Table S1). Nearly all Isolates predicted to produce an ESBL were non-susceptible to ampicillin/sulbactam (100%), aztreonam (94%), cefazolin (100%), cefepime (94%), ceftriaxone (100%) (Table S1). Only one isolate, KPN103, had considerable beta-lactam resistance but lacked an ESBL gene; however, this isolate did contain the gene encoding the non-ESBL beta-lactamase, *bla_TEM-1D_* (Table S1).

Of the 104 *K. pneumoniae* isolates, 34 (32.6%) were non-susceptible to the combination of trimethoprim and sulfamethoxazole (TMP/SMX) (Fig S3). Of these isolates 31 (29.8%) contained both the *dfrA* and *sul1/2/3* genes, which encode resistance to TMP/SMX. (Table S1). Two resistant isolates contained only *dfrA* and one contained only *sul2* (Table S1).

Eighteen isolates (17.3%) were resistant to ciprofloxacin (Fig S3). Each of these isolates contained either *qnrB* gene, or *a* mutation in the *gyrA* gene (Table S1).

Within the collection, non-susceptibility rates to tobramycin, gentamicin, and amikacin were 17.3%, 10.5%, and 1.9%, respectively (Fig S3). The overall prevalence of aminoglycoside modifying enzymes (AMEs) was 31.7% (n=33) (Table S1). The most common AMEs were streptomycin resistance genes *strA* and *strB* (n=22) and the streptomycin/spectinomycin adenyltransferase *aadA2* (n=10); however, isolates containing only these AMEs were sensitive to tobramycin, gentamicin, and amikacin (Table S1). The acetyltransferases *aac(6’)-Ib-cr.V2* (n=7), *aac(3)-IId* (n=2), and *aac(3)- IIa* (n=2), and the phosphotransferases *aph(3)-Ia.v1 (n=5) and aph(3’)-Ia (n=2)* were associated with nonsusceptibility to gentamicin and tobramycin (Table S1).

Collectively, 17 (16.3%) of the 104 isolates were MDR including 12 (11.5%) that were extensively drug-resistant (XDR) (Fig 2). These findings indicate the MDR bacteria comprise a substantial proportion of *K. pneumoniae* causing bloodstream infections at NMH and suggest that many strains carry mobile elements with multiple resistance genes.

### Virulence Determinants

We next searched the *Klebsiella* genomes for genes associated with enhanced virulence. The siderophore yersiniabactin promotes pulmonary infections in animal models, is associated with more severe disease in human infections, and is produced by hvKP strains (4, 39, 51, 52). We identified yersiniabactin biosynthesis loci (*ybt*) in 37.5% (n=39) of 104 NMH bloodstream isolates (Fig 2 & Table 2). Eleven unique alleles of *ybt* were detected, the most common being *ybt* 10 (n=12), *ybt* 17 (n=11) and plasmid-borne *ybt* 4 (n=4) (Table S1). *ybt* and the colibactin biosynthesis locus, *clb*, are commonly found together on an integrative conjugative element, *ICEKp10* (18). We identified 12 (11.5%) isolates that contained *ICEKp10* representing five STs (ST258, ST29, ST66, ST15, and ST240) (Table S1). Although *ICEKp10* is a marker for the globally distributed CG23-I sub-lineage, within the United States, *ICEKp10* is a strong marker for the epidemic ST258 lineage (18, 53).

In addition to *ybt* and *clb*, we screened our collection for the following previously published diagnostic biomarkers of hvKP: *iucA, iroB, rmpA, rmpA2,* and the hmv phenotype. *iucA* was identified in 5 (4.8%), *iroB* in 4 (3.8%), *rmpA* in 4 (3.8%), and *rmpA2* in 2 (1.9%) (Fig 2 & Table 2). These virulence genes tended to be clustered within the same few strains. Four isolates contained *iucA, iroB,* and *rmpA* (Table 1). Two of these four (KPN8 and KPN115) were ST23, a sequence type commonly associated with hvKP, and the other two (KPN49 and KPN165) were ST66 and ST380, respectively. A single isolate, KPN23, was ST881 and contained *iucA* but not *iroB* or *rmpA*. (Fig 2 & Table 2). Three distinct *iucA* alleles were identified: *iuc*1 (n=2, ST23), *iuc2* (n=2, ST66 & ST380), and *iuc3* (n=1, ST881) (Table 1). Although two strains contained the *rmpA2* gene, both had a frameshift mutation in a previously published homopolymer region of *rmpA2* (54). All *iuc+* isolates were sensitive to every antibiotic tested, except ampicillin (Table S1). The collection was screened for hmv capsules by both string test and centrifugation. A total of 10 (9.6%) and 18 (17.3%) isolates, respectively, were positive for hypermucoviscosity by these two tests (Table 2), including all four of the isolates containing *iucA, iroB,* and *rmpA* but not the isolate containing only *iucA* (Table 1, Table S1). These data confirm previous reports that the string test is less sensitive than centrifugation for measuring hypermucoviscosity (48, 55). However, all *rmpA*+ isolates were positive by both string and centrifugation tests (Table S1). In addition, isolates were tested for tellurite resistance, a phenotype commonly encoded by pK2044 plasmids (Fig 1). Five isolates (4.8%) grew on tellurite, including the two hmv ST23 isolates that were positive for *rmpA, rmpA2*, *iro1* and *iuc1 alleles* (Table 2). These results suggested that these two isolates contained pK2044-like plasmids, while the two isolates containing *iro2, iuc2* and *rmpA* contained KP52.145pII-like plasmids. Collectively, these findings indicate that clinical isolates with genomic features of hvKP are circulating at NMH.

**Table 1.**
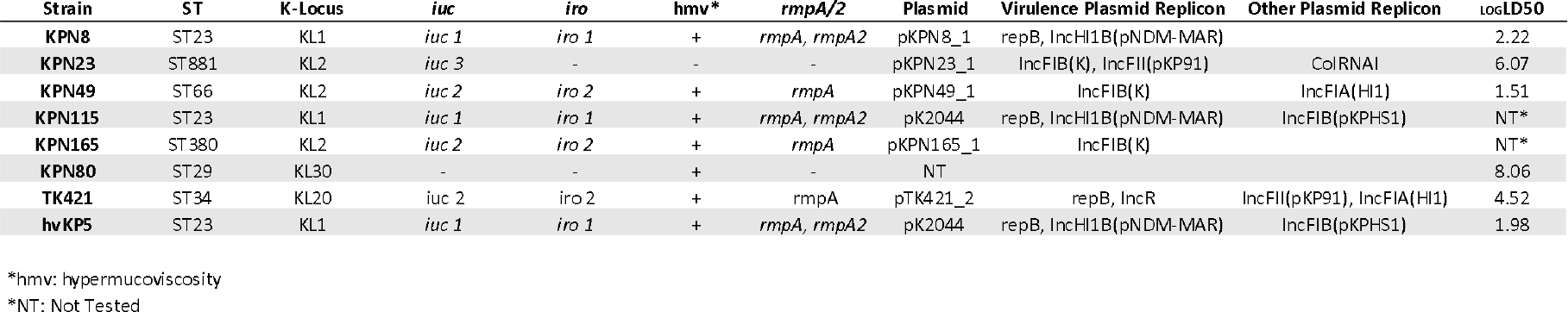
Genomic Characteristics of bloodstream isolates containing hvKp pathogenicity loci.

**Table 2.** Virulence factors identified in NMH bloodstream isolates.

| Isolates n = 104 | Positive<br>n(%) |
| --- | --- |
| *iroB* | 4 (3.8) |
| *iucA* | 5 (4.2) |
| *rmpA* | 4 (3.8) |
| *rmpA2* | 2 (1.9) |
| *ybt* | 39 (37.5) |
| *clb* | 12 (11.5) |
| *iroB/iucA/rmpA* | 4 (4.2) |
| String Test | 10 (9.6) |
| Centrifugation Test | 18 (17.3) |
| Tellurite Resistance | 5 (4.8) |

### Analysis of Plasmid Content

Next, we sought to characterize the plasmids harbored by the five isolates containing salmochelin and/or aerobactin biosynthesis genes. To define plasmids sequences, we performed Nanopore long read sequencing on these isolates. Hybrid assembly with the Illumina short reads of the genomes revealed that each of these isolates harbored a large plasmid with some hvKP pathogenicity loci (Table 1). Two isolates (KPN8 and KPN115) contained plasmids with near complete alignment with pK2044, including plasmid replicons, *repB* and *IncHI1B* and pathogenicity loci, PAL-1 and PAL-2 (Fig 3 and Table 1). KPN165 harbored an *IncFIB(K)* plasmid with close alignments with KP52.145pII but not pK2044 (Fig 4A and S5). Similar to KP52.145pII, this plasmid contained a version of PAL-1 with *iucABCD* and *iutA* but lacking *rmpA2* (Fig 1 and 4A). Interestingly, KPN49 contained two plasmids: one (pKPN49_1) with an *IncFIB(K)* replicon and homology to KP52.145pII and a second (pKPN49_2) with an *IncFIA(HI1)* replicon and homology to KP52.145pI (Fig 4A and Table 1). However, the KP52.145pII-like plasmids of KPN49 and KPN165 were ∼40 kB larger than KP52.145pII. Analysis of this 40 kB sequence revealed that it contained a *tra* conjugation locus with high sequence homology with KP52.145pI, a second plasmid contained by reference hvKP strain KP52.145 (Fig 4B) (34, 56). We also identified a similar chimeric hypervirulence plasmid (99.0% identity) in a previously published ST380 strain, hvKP4 (Fig 4B) (57, 58). The fifth isolate, KPN23, carried on a two-replicon plasmid (*IncFIB(K)* and *IncFII(pKP91)*) with aerobactin biosynthetic genes and putative conjugative genes. This plasmid did not resemble known hypervirulence plasmids but was highly similar to a plasmid harbored by a swine isolate KPCTRSRTH01_p2 collected in Thailand in 2016 (Fig 5A & Table 1). For comparison, we included in the analysis pTK421_2, a unique hvKP plasmid we had previously identified from a bloodstream isolate collected at NMH in 2013 (35). pTK421_2 contains PAL-1, PAL-2, and conjugation genes (Fig 1). This plasmid was also distinct from pK2044 or KP52.145pII (Fig 4A & S5). We therefore aligned pTK421_2 to pKPN23_1 but found that these plasmids shared relatively little alignment beyond PAL-1 (Fig 5B). Overall, among the 104 *K. pneumoniae* bloodstream isolates, we identified four isolates that contained two distinct hypervirulence plasmids and a fifth isolate that contained a plasmid containing aerobactin biosynthetic genes.

**Figure 3.**
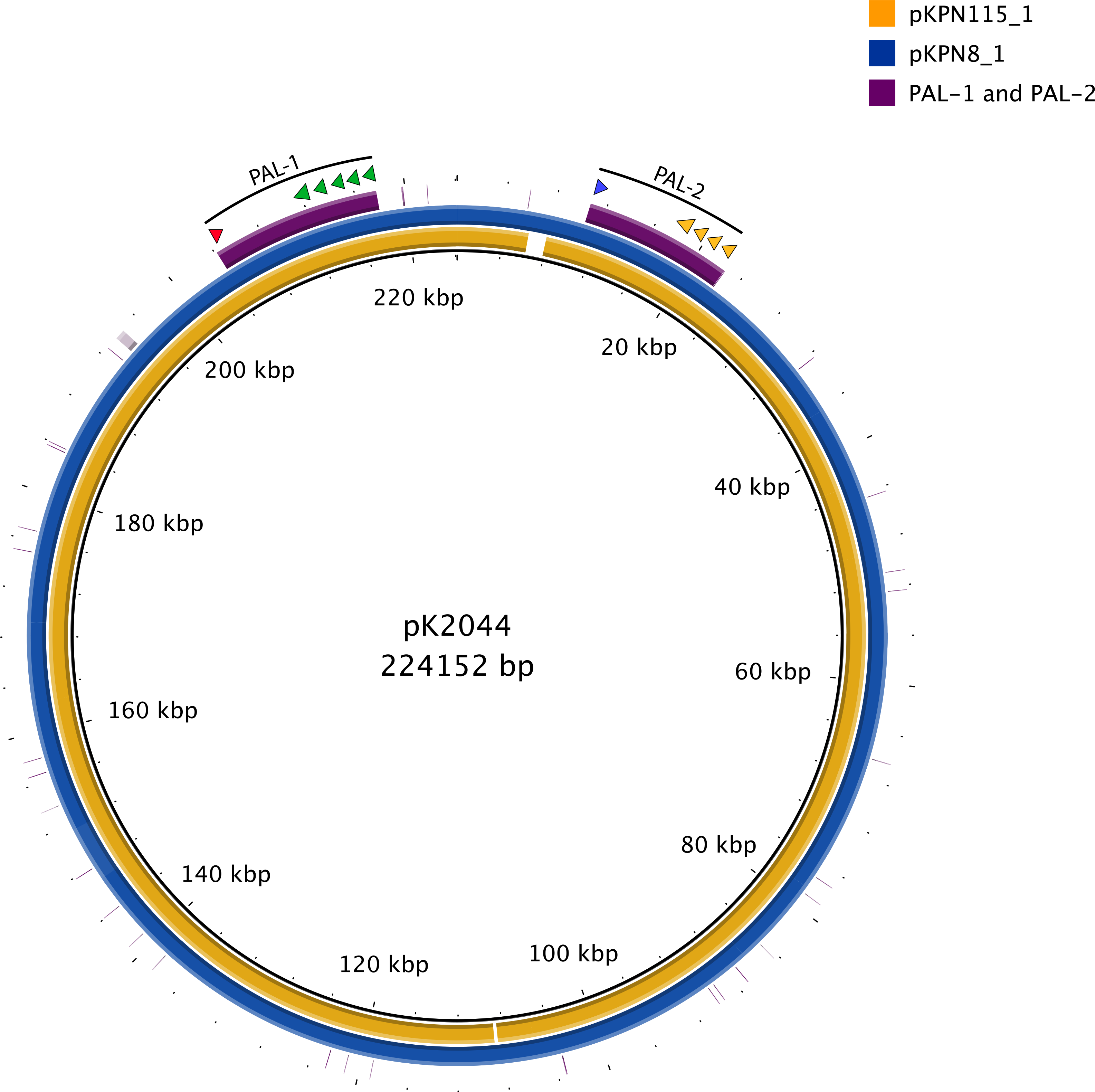
The bloodstream isolates KPN115 and KPN8 harbor pK2044-like plasmids. pKPN115_1 and pKPN8_1 sequences were aligned to pK2044 using blast ring image generator (BRIG). A sequence identity threshold of 85% was used. Aerobactin biosynthesis genes are indicated with green arrows, salmochelin with orange arrows, *rmpA* with a blue arrow, and *rmpA2* with a red arrow.

**Figure 4.**
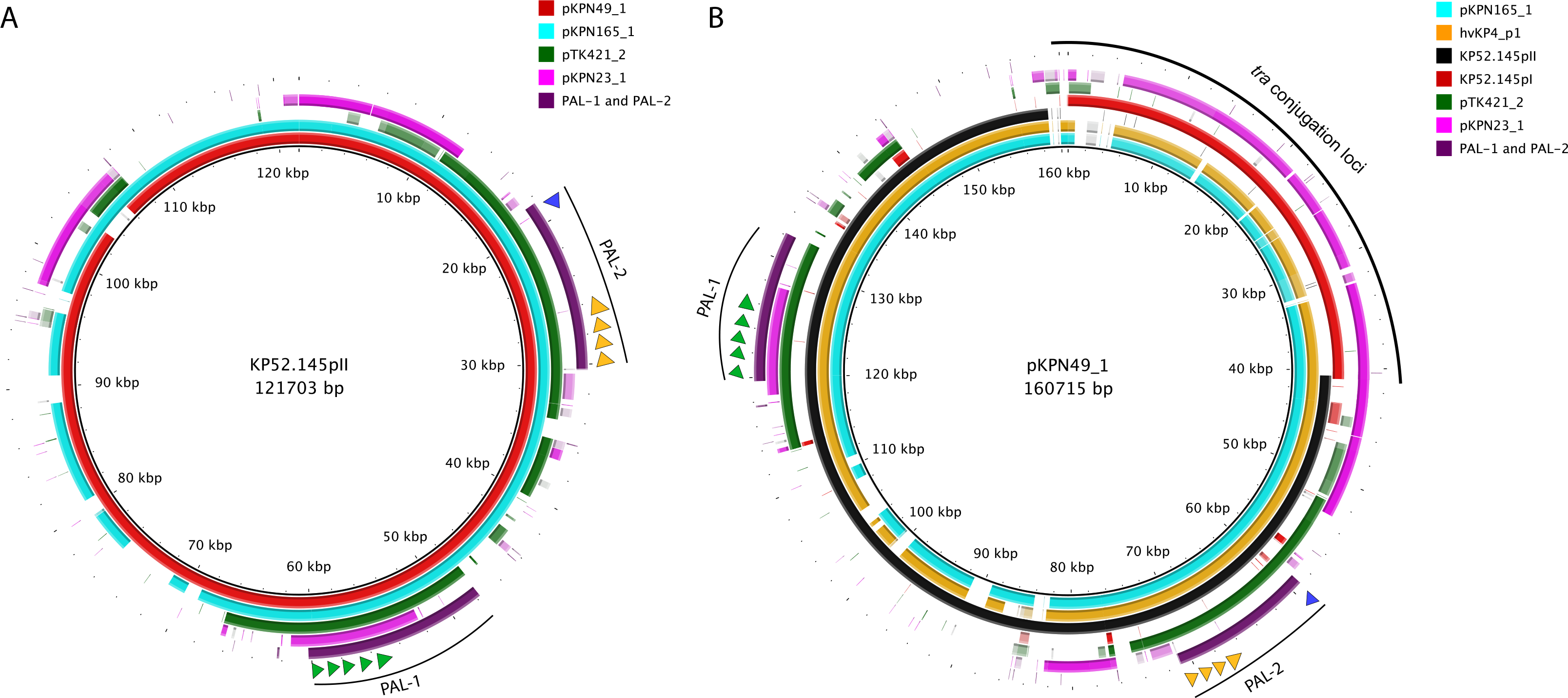
The bloodstream isolates KPN49 and KPN165 harbor KP52.145pII-like plasmids. **A.** The indicated plasmids sequences were aligned to KP52.145pII using blast ring image generator (BRIG). **B.** The indicated plasmids sequences were aligned to pKPN49_1 using blast ring image generator (BRIG). A sequence identity threshold of 85% was used. Aerobactin biosynthesis genes are indicated with green arrows, salmochelin with orange arrows, and *rmpA* with a blue arrow.

**Figure 5.**
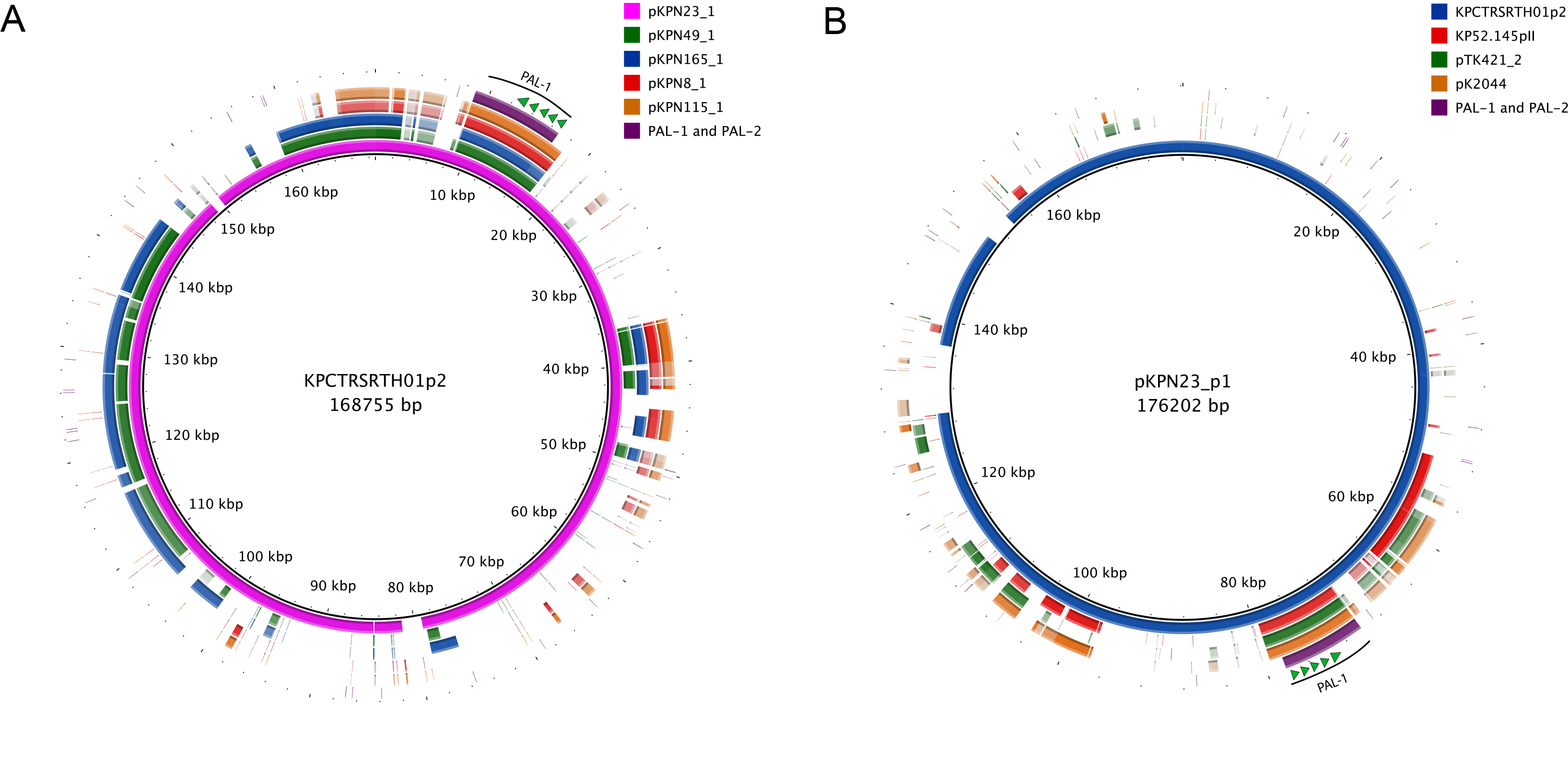
The bloodstream isolate KPN23 harbors a unique plasmid containing aerobactin biosynthesis genes. The indicated plasmids were aligned to that of KPCTRSRTH01_p2. **(A)** or pK2044, KP52.145pII, and pTK421_2 **(B)** were aligned with pKpN23_1 using blast ring image generator (BRIG). A sequence identity threshold of 85% was used. Aerobactin biosynthesis genes are indicated with green arrows. Aerobactin biosynthesis genes are indicated with green arrows.

### Phylogenetic analysis of bloodstream isolates containing hvKP pathogenicity loci

To examine the phylogenetic relationships of the five NMH bloodstream isolates containing aerobactin biosynthesis genes, we generated a core genome phylogenetic tree of these isolates along with previously published hvKP isolates. These results confirmed previous findings that KL1 hvKP isolates tend to be phylogenetically clustered, whereas KL2 hvKP isolates tend to be more dispersed (Fig S6). Neither ST23-K1 isolate is part of a previously described and globally distributed CG23-I clade and, as such, neither contained the combination of yersiniabactin and colibactin biosynthetics genes observed in *ICEKp10* (Fig S6) (18). KPN49 is part of a rarely described ST66-K2 lineage consistent with the recent global re-emergence of this clone (Fig S6) (56, 59, 60).

Although KPN165 and KPN49 both harbor KL2 capsule loci and very similar KP52.145pII-like plasmids (99.6% identify) containing additional putative conjugation loci, they are quite divergent genetically (Fig 4B, S6, & Table 1). These findings suggest that this plasmid, which contains a number of virulence genes, has spread by horizontal transmission. Together, these results suggest that nonclonal strains with features of hypervirulent *K. pneumoniae* are circulating at NMH.

### Pathogenesis in a murine model of pneumonia

Studies to date suggest that hvKP strains share the common feature of being highly virulent in mouse models of infection. To more definitively determine whether the isolates with features of hvKP were indeed hypervirulent, we measured the virulence of representative isolates in a mouse pneumonia model. To determine the level of virulence of the isolates containing characteristics of hvKP, 50% lethal dose (LD_50_) measurements were made. For reference, we first determined the virulence of a published hvKP isolate, hvKP5, which is an ST23 KL1 strain that harbors *iuc1, iro1, rmpA,* a disrupted allele of *rmpA2,* and is hmv (44). As expected, hvKP5 was extremely virulent and had a LD_50_ of 10^2.0^ CFU. In this model hvKP5 was more virulent than previously published hvKP isolates, hvKP1 and hvKP4, which have an LD_50_ of 10^3.2^ and 10^4.6^ CFU, respectively (Fig 6A) (57). From our collection, we selected KPN8 and KPN49 as representative pK2044 KP52.145pII isolates, respectively. These isolates had LD_50_ values of 10^2.2^ and 10^1.5^ CFU, respectively, indicating high levels of virulence (Fig 6C & 6E). In comparison, KPN23, which is non-hmv and has an *iuc3* virulence plasmid lacking other hvKP virulence genes, had an intermediate LD_50_ value of 10^6.1^ CFU (Fig 6D). We also included in the analysis TK421, the hvKP-like strain that we had previously identified that contained *iuc2, iro2, rmpA,* and was hmv (Table 1). TK421 had an LD_50_ of 10^4.5^ CFU (Fig 6B). Finally, as a control we tested KPN80, an hmv blood stream isolate from our collection lacking *iuc, iro, rmpA,* and *rmpA2* genes (Table 1) (35, 44). KPN80 had an LD_50_ value of 10^8.1^ CFU which is consistent with previously published LD_50_ values for cKP (Fig 6F) (57). Collectively, these data confirm that KPN8, KPN49, and TK421 are hvKP strains and suggest that KPN115 and KPN165 are also hvKP. KPN23 had an intermediate pathogenic phenotype and is more virulent than a cKP strain but less virulent than typical hvKP strains. In addition, these findings indicate that there is a spectrum of virulence among hvKP isolates. These data suggest that multiple factors are necessary for full virulence and that *iuc* genes or a hmv phenotype alone is not sufficient to define an isolate as hypervirulent.

**Figure 6.**
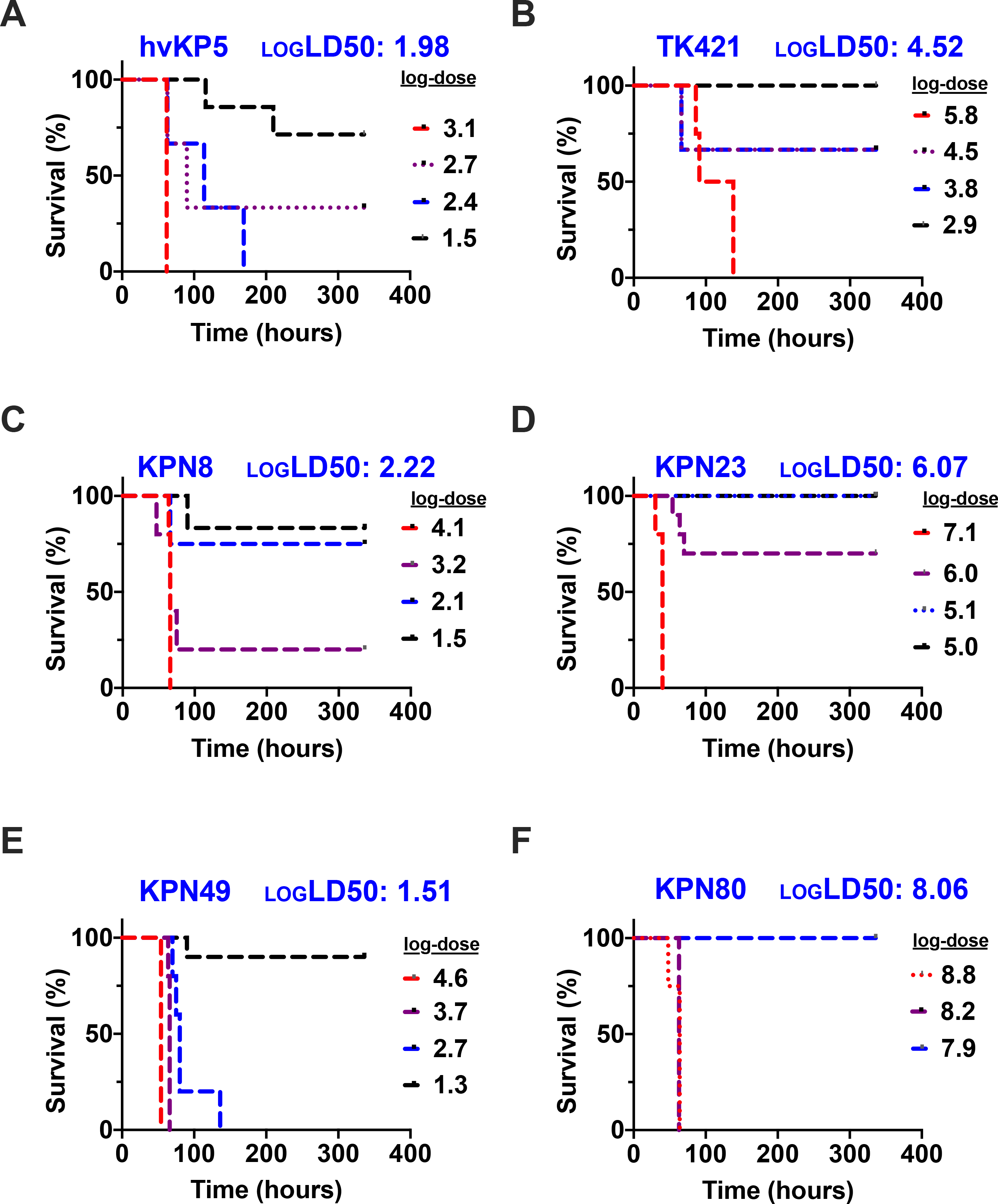
NMH Isolates containing hvKp pathogenicity loci were extremely virulent in a murine model of pneumonia. C57BL/6 mice were infected by nasal aspiration with the indicated doses of hvKP5 **(A)**, TK421 **(B)**, KPN8 **(C)**, KPN23 **(D)**, KPN49 **(E)**, and KPN80 **(F**). Genotypes, phenotypes, and calculated LD50s are listed in Table 1.

### Infections caused by hvKP strains

Since KPN8, KPN49, KPN115, and KPN165 had microbiological, genetic, and virulence properties of hvKP, we next examined the clinical context of the infections they caused. None of the patients infected by these strains had known travel history to areas where hvKP is endemic. Three of the four patients had infections that were community-acquired. Two infections led to hepatic abscesses, and one infection resulted in a lung abscess (Table 3). Two infections occurred in patients with diabetes mellitus, a known risk factor for hvKP (Table 3) (61-63). None of the patients had other common manifestations of hvKP infections, such as endophthalmitis, meningitis, or necrotizing fasciitis. We also looked at the infections cause by KPN23, KPN80, and TK421. The patient infected with TK421, a slightly less virulent strain with features of hvKP, had a history of diabetes mellitus and presented with a community-acquired liver abscess, although this patient had predisposing conditions including a previous Whipple procedure and a history of prior liver abscesses. The patient infected with KPN23, the *iuc3*+, non-hmv strain with somewhat lower virulence, did not have features of or risk factors for hvKP infection. Likewise, the patient infected with KPN80, the hmv isolate that lacked hvKP-associated genes, also did not have features of or risk factors for hvKP infection. Although the very small number of patients prevents conclusions from being drawn, the characteristics of these infections support the assertion that hvKP strains are circulating at NMH.

**Table 3.** Clinical Charac eris ics o bloods ream isola es con aining hvKp° pa hogenici y loci.

| Strain | Age (Years) | Type of Infection | Community-Acquired Infection | Medical Comorbidities |
| --- | --- | --- | --- | --- |
| KPN8 | 67 | Bloodstream only | No | Type II Diabetes, Myocardial Infarction, Peripheral Vascular Disease |
| KPN23 | 50 | Bloodstream only | No | Pancreatic Adenocarcinoma |
| KPN49 | 53 | Pneumonia/lung abscess | Yes | HIV |
| KPN115 | 73 | Hepatic Abscess | Yes | Type II Diabetes, Colon Adenocarcinoma |
| KPN165 | 50 | Hepatic Abscess | Yes | Cholangiocarcinoma |
| KPN80 | 68 | Pneumonia | Yes | Cirrhosis |
| TK421 | 65 | Hepatic Abscess | Yes | Type II Diabetes, Pancreatic Adenocarcinoma |

\* Hypervirulent *Klebsiella pneumoniae*

## Discussion

*K. pneumoniae* bloodstream isolates at NMH had significant diversity in sequence type, capsule type, antimicrobial resistance genes, and virulence factors. In total, there were 75 distinct sequence types in our collection, demonstrating that a broad spectrum of the *K. pneumoniae* species has the potential to cause invasive infections. Nearly all of these isolates (n=100, 96%) were cKP (i.e. lacked genetic features of hvKP, and a substantial proportion of these were MDR or XDR (n=17, 16.3% and n=12, 11.5%, respectively) and 9 (8.7%) were resistant to carbapenems. A substantial number of the isolates belonged to the established global high-risk clones including ST258, ST45, ST23, ST15, ST66, and ST147. However, 4 (3.8%) of the isolates had genetic, phenotypic, and virulence features consistent with hvKP, indicating that hvKP strains are circulating in the U.S. community. These four isolates belonged to the established hvKP ST23 and ST380 lineages and the reemerging ST66 lineage. These findings indicate that antibiotic-resistant and high-risk clones comprise a substantial number of *K. pneumoniae* causing bloodstream infections at NMH but that hvKP strains are also present.

In MDR cKP infections, delay in administration of active antimicrobial therapy may lead to poor outcomes (64, 65), and it therefore important to know the prevalence of these infections. Two large studies have recently evaluated the molecular characteristics of either extended spectrum beta-lactamase resistant or carbapenem resistant isolates from U.S. hospitalized patients (from 2011-2017) and ST258 and ST307 are found at exceptionally high rates, 26.7-64.4% and 7.4-35.7%, respectively (66, 67). A recent study from our group showed that at NMH 74.1% of carbapenem resistant isolates are ST258 (68). Among bloodstream isolates at NMH, 17 (16.3%) isolates were resistant to three or more classes of antibiotics and considered MDR-cKP, 12 (11.5%) isolates were ESBL producing (12%). Comparatively, ESBL prevalence among collections from Asia ranges from 12-79% (69-73). Carbapenamases were found in 9% of isolates, with most strains carrying *bla_KPC-3_* and one strain carrying *bla_NDM-1_*. As expected, ST258 (n=6, 5.7%) and ST307 (n=2, 1.9%) were both identified in our collection.

The invasive nature of hvKP strains allow them to cause unusual infections such as PLA, endogenous endophthalmitis, necrotizing fasciitis, and meningitis associated with clinical implications distinct from those of the more common cKP strains (33). For example, 9-18% of PLA patients have a recurrence of their infection (74, 75), suggesting that prolonged treatment courses may be necessary. In one study, 10 of 14 patients with endogenous endophthalmitis progressed to blindness (76), underscoring that these infections are medical emergencies. hvKP strains are prone to disseminate, and culture of an hvKP isolate from, for example, a liver abscess may prompt a search for other foci of metastatic infection. It therefore can be argued that hvKP isolates should be identified as such by hospital laboratories and that the prevalence of these isolates should be studied. Our finding that 3.8% of *K. pneumoniae* bloodstream isolates at NMH were hvKP agrees with two published U.S. surveillance studies, which describe the hvKp prevalence as 3.7%-6.3% of *K. pneumoniae* infections in two U.S. cities. In 2010, 64 *K. pneumoniae* strains were analyzed from two hospitals in the greater Houston area (25). They identified four strains (6.3%) containing hvKp pathogenicity-associated genes, based on the presence of *rmpA* and K1 capsule type by PCR amplification. In 2018, a larger study in New York City characterized *K. pneumoniae* isolates from 462 patients (24). This study identified 17 (3.7%) isolates by PCR amplification that contained the hvKp biomarkers *rmpA* and *iucA*. In contrast, a 2013 report from China, where hvKP is endemic, described an hvKP prevalence of 37.8%, (defined by the presence of the *iuc* gene) (77). A recent genomic surveillance study of seven countries in South and Southeast Asia analyzed the sequences of 365 *K. pneumoniae* bloodstream isolates collected from 2010-2017 (78). Sequencing revealed that 26% (95/365) were *iuc*+, 17% (63/365) were *iuc*+ and either *rmpA*+ or *rmpA2*+, and an additional 0.8% (3/365) were *iro*+ and *rmpA*+. Alarmingly, of the *iuc*+ isolates 13.7% (13/95) contained carbapenamase genes, but only 1 was positive for multiple hvKp factors (*iuc*, *rmpA/2*) and a carbapenamase gene. Thus far, antimicrobial-resistant hvKp infections are rare in the United States, and none of the *iuc*+ isolates identified at NMH contained accessory antimicrobial resistance genes. However, antimicrobial-resistant hvKp infections are on the rise in Asia, likely due to the increase in hospital-acquired hvKp infections (79-81). Our results together with these reports indicate call attention to the need for further studies evaluating the prevalence and clinical implications of hvKP infections in the United States and other countries outside of Asia.

Since hvKp pathogenicity loci are normally encoded on large plasmids, we performed long-read sequencing of the isolates that contained *iucA*. Four of these isolates harbored a plasmid similar to one of the two previously described hvKp virulence plasmids, pK2044 or KP52.145pII (28-30). The plasmids from these four isolates contained multiple hvKP-associated genetic loci (*iuc, iro,* and *rmpA*), and the isolates that harbored them had hmv phenotypes and LD_50_ values consistent with hvKp (Fig 6 and Table 1). Among these, three isolates (KPN49, KPN115, and KPN165) caused severe community-acquired infections with abscess formation (2 hepatic and 1 lung). Taken together, these data indicate that these four isolates are indeed hvKP and that multiple strains of hvKp with distinct plasmids are circulating at NMH.

A closer examination of the isolates and plasmids provided several additional insights into hvKP. For example, the KP52.145pII-like plasmids of KPN49 and KPN165 contained a 40 kb insert containing putative conjugation genes, suggesting that these plasmids, unlike typical hypervirulence plasmids, were capable of transmission. Indeed, the two isolates that contained these plasmids, KPN49 and KPN165, were phylogenetically distinct (ST66 and ST380, Fig S6), consistent with horizontal dissemination of this plasmid. We identified a number of isolates that were hmv but did not have other features of hvKP. One of these isolates, KPN80, was tested in the mouse model and was found to have a very low level of virulence. These findings agree with previous reports indicating that hypermucoviscosity does not accurately predict hvKP (48, 55). In addition to the four hvKP isolates, we identified one additional isolate, KPN23, that harbored a unique plasmid containing the aerobactin biosynthesis locus but not other hvKP-associated virulence genes. This isolate was not hmv, and it caused a typical cKP disease: a hospital-acquired infection in a patient with pancreatic cancer that was not associated with abscess formation. However, it was associated with a level of virulence in the mouse model that was intermediate between hvKP and cKP. These findings suggest that *K. pneumoniae* strains may not fall neatly into either the cKP or the hvKP classification but may have intermediate phenotypes.

## Conclusions

In this study, *Klebsiella spp*. bloodstream isolates at NMH were collected consecutively from 2015-2017. Whole genome sequencing revealed that these isolates are diverse in sequence type, capsule type, and virulence gene content. 30.7% of isolates were high-risk clones, predominantly ST258 and ST45, whereas 3.8% of isolates were hvKP belonging to ST23, ST66, and ST380.

## Materials and Methods

### Bacterial isolates and growth conditions

The isolates used in this study were from consecutive blood cultures collected between April 2015-April 2017 with growth designated as *K. pneumoniae* by the NMH Clinical Microbiology Laboratory using a VITEK-2 platform (82). For the sub-analysis of isolates determined to be *K. pneumoniae* by sequencing, only the first isolate cultured from each unique infection was included. For experiments, bacteria were grown at 37°C in lysogeny broth (LB) or on LB agar plates.

### Whole-Genome Sequencing

Genomic DNA was prepared from a single colony cultured overnight at 37°C in LB using the Maxwell 16 system (Promega Corp., Madison, WI). Libraries for Illumina sequencing were prepared using either Nextera XT (Illumina, Inc., San Diego, CA) or Seqwell (Seqwell, Massachusetts) library kits and sequenced on an Illumina MiSeq or NextSeq 500 instrument to generate paired-end 300 bp or 150 bp reads (Table S3). Reads were trimmed using Trimmomatic (v0.36) and *de novo* assembly was performed using SPAdes version 3.9.1 (83, 84). Nanopore sequencing was performed as described previously (85, 86). Briefly, libraries were prepared from genomic DNA using the ligation sequencing kit (SQK-LSK109, Oxford Nanopore, UK). Libraries were sequenced on the MinION using a FLO-MIN106 flow cell and base calling and demultiplexing of sequence reads was performed using Guppy v3.4.5 (87). Hybrid assembly and circularization of Nanopore and Illumina reads were performed using Flye v2.9. Flye assembly information is included in Table S4. Nanopore sequencing errors were corrected by aligning Illumina reads to the assembly using BWA v0.7.17 and correcting the errors with serial rounds of Pilon v1.23. Annotation was performed using the NCBI Prokaryotic Genome Annotation Pipeline (88).

### Hypermucoviscosity testing

The string test was performed as described previously (89). Isolates were grown overnight at 37°C on LB agar. A single colony was lifted with a loop to evaluate the formation of a viscous string between the loop and the colony. A positive string test was defined as a string length ≥ 5 mm. A centrifugation assay was also used to assess hypermucoviscosity. Centrifugation of overnight bacterial cultures (5 mL LB). was performed at 3,220 x g (RCF) for 10 minutes (55). Hypermucoviscous isolates were identified qualitatively by the persistence of turbidity.

### Tellurite resistance testing

For screening of tellurite resistance, *K. pneumoniae* was grown at 37°C on LB agar plates supplemented with 3 μg/mL potassium tellurite (Sigma-Aldrich, USA).

### Phylogenetic analysis

FastTree 2 was used to estimate a maximum-likelihood phylogenetic tree (90). Core SNPs were determined by aligning raw Illumina reads to the reference strain (NTUH- K2044) using bwa-0.7.15 and filtering them to 95% core. The phylogenetic tree was visualized and annotated using iTOL (v4)(91).

### Molecular typing and identification of virulence genes

Assembled whole genome sequences were analyzed with the bioinformatics tools *Kleborate* v2.1 and *Kaptive* v0.7.3 to evaluate multilocus sequence type, capsule locus typing, antimicrobial resistance genes, and virulence gene content (31, 92). Plasmids identified by whole genome sequencing were aligned using BLAST Ring Image generator (BRIG) with the alignment threshold set at 85% identity (93). Plasmid replicons were identified using the PlasmidFinder v2.1 online database (94).

### Animal Studies

Anesthetized 6- to 8-week-old C57BL/6 female mice purchased from Jackson Labs were infected intranasally as described previously (48). Briefly, sets of mice were infected with different doses and monitored for two weeks post-infection for pre-lethal illness. Strain dosing, deaths, and total mice inoculated are included in Table S2. LD_50_ values were determined in R, with the *drc* package (95). To minimize pain and distress in the animals, mice were euthanized when they reached predetermined endpoints of >20% weight loss, abnormal respiratory rate, or a hunched posture with minimal activity. All procedures were performed in accordance with the guidelines approved by the Northwestern University Animal Care and Use Committee as described in protocol IS00002172.

## Statistical Analysis

Statistical analysis was performed using GraphPad Prism. Student T-test and analysis of variance (ANOVA) followed by the Bonferroni’s correction for multiple comparisons were performed for parametric variables. For non-parametric variables, the Mann-Whitney U test was used. For comparison of survival curves, the Mantel-Cox log rank test was used. Simpson Diversity index was determined in R using the package ‘vegan’.

## Declarations

### Ethics, consent and permissions

#### Patient Chart Review

The following information was extracted retrospectively from the electronic medical records of patients from which the blood cultures were obtained: antibiotic susceptibility (as determined by the NMH Clinical Microbiology Laboratory using a VITEK-2 platform), age, type of infection, medical comorbidities, travel history, and community vs. hospital acquisition of the *K. pneumoniae* infection (i.e. onset prior to 48 hr of hospitalization or after 48 hr of hospitalization). Clinical breakpoints used to categorize isolates as resistant, intermediate, or susceptible are described for each antibiotic in Table S1. All data were collected as part of routine diagnostic care and all samples and data have been de-identified and pose no more than minimal risk to subjects. This study was approved and granted a waiver of the consent process by the Northwestern Institutional Review Board under protocol STU00211033.

### Consent to publish

Not applicable

### Availability of data and materials

The whole-genome assemblies have been deposited at GenBank under BioProject number PRJNA788509. Assembly accession numbers are included in Table S3. MIC and *Kleborate* data are included in Table S1.

### Authors’ contributions

TK and AH designed the study. TK, SN, BC, SG, MLC SM, and NK performed the phenotypic experiments. TK and EO performed the computational analyses. RM and BC analyzed and interpreted the patient data. FK and CQ collected and identified the isolates as *Klebsiella spp*. TK, SN, SM, EO, and AR analyzed the data. TK and AH wrote the paper, and all other authors contributed to the writing. All authors read and approved the final version of the manuscript.

## Supplemental Figure Legends

**Figure S1. Population structure of *Klebsiella* spp. bloodstream isolates.** Maximum likelihood phylogenetic tree generated from core genome SNP loci in 140 *Klebsiella* spp. bloodstream isolates. The tree has a truncated outlier branch for *Klebsiella oxytoca* (yellow). The scale bars represent genetic distances.

**Figure S2. NMH *K. pneumoniae* bloodstream isolates are highly diverse in ST, KL, and O antigen.** Numbers of genomes with each corresponding ST **(A)**, KL **(B)**, or O-antigen type **(C)** were determined using *Kleborate* and *Kaptive*.

**Figure S3. Antimicrobial resistance phenotypes of *K. pneumoniae* bloodstream isolates.** The percentage of isolates resistant (solid bars) or intermediately susceptible (cross-hatched bars) to the indicated antibiotics are shown. Antibiotic classes are grouped by color.

**Figure S4. Beta-lactam resistant isolates contain a variety of extended-spectrum beta-lactamase or carbapenamase genes.** Numbers of genomes containing the indicated beta-lactamase genes present in isolates resistant to ceftriaxone **(A)**, meropenem **(B)**, amp/sulbactam **(C)**, aztreonam **(D)**, cefazolin **(E)**, or cefepime **(F)** are plotted. “-“ indicates strains without an extended-spectrum beta-lactamase.

**Figure S5. pKPN49_2, pKPN165_1, pTK421_2, and pKPN23_1 have little similarity to pK2044.** DNA sequences of these plasmids were aligned to pK2044 using blast ring image generator (BRIG). A sequence identity threshold of 85% was used. Aerobactin biosynthesis genes are indicated with green arrows, salmochelin with orange arrows, *rmpA* with a blue arrow, and *rmpA2* with a red arrow.

**Figure S6. Core genome phylogenetic tree of United States hypervirulent *Klebsiella pneumoniae* isolates**. A maximum likelihood phylogenetic tree was generated from core genome SNP loci in hvKP isolates from United States hospitals and select global reference isolates. Sequence types are indicated. The presence of virulence factors is listed next to each isolate: *ybt* = yersiniabactin biosynthesis loci, *clb* = colibactin biosynthesis loci, *rmpADC* = mucoid regulator operon, *rmpA2* = regulator of mucoid phenotype 2, *iuc* = aerobactin biosynthesis genes, *iro* = salmochelin biosynthesis genes. The scale bars represent genetic distances. KPPR1, KP52.145, SB5881, K180005, NCTC9494, NTUH-K2044, RJF999, and ED2 are global isolates used as a reference.

