## Supplementary figures and images for "Genomic surveillance for multidrug-resistant or hypervirulent *Klebsiella pneumoniae* among bloodstream isolates at a United States academic medical center"

### Figure S1

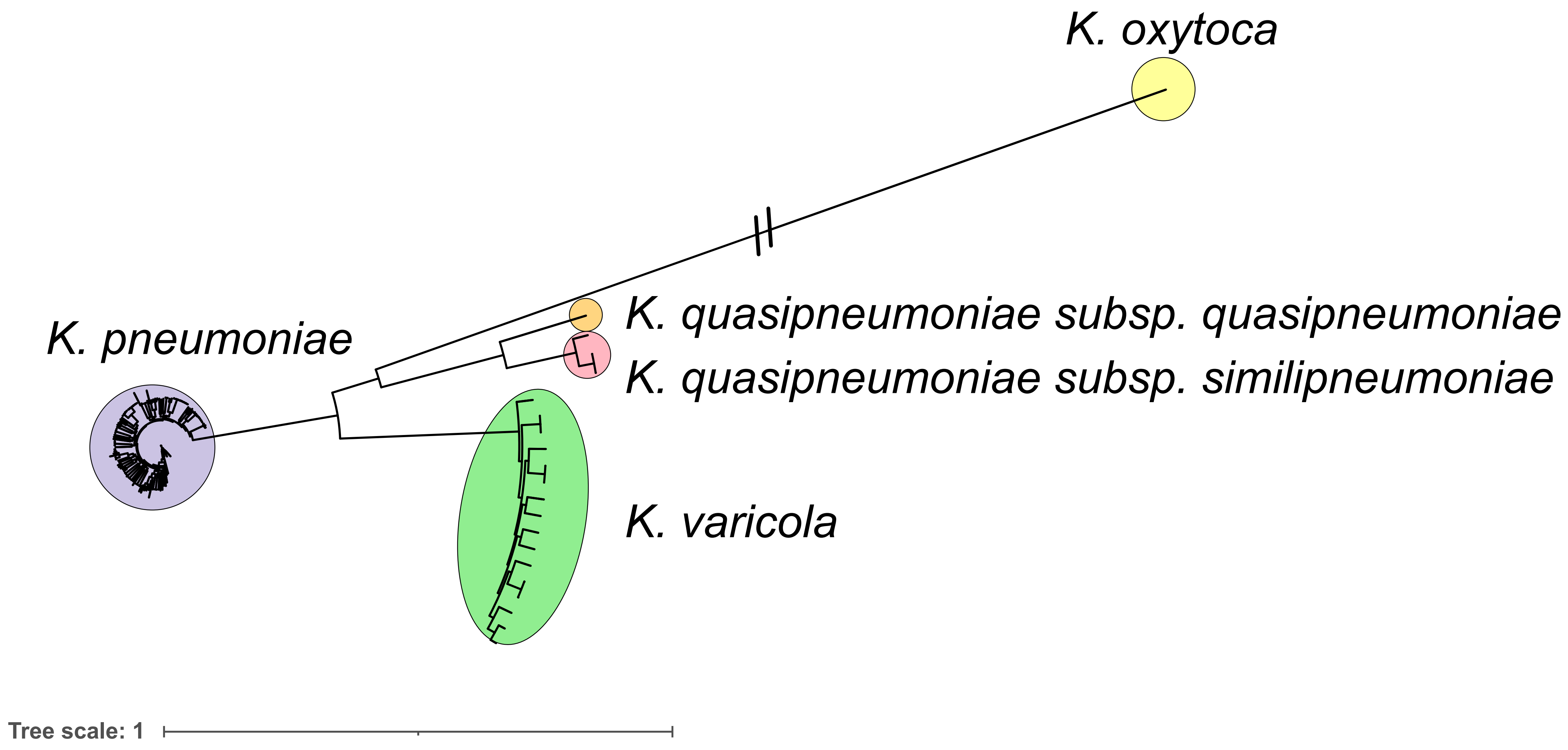

### Figure S2

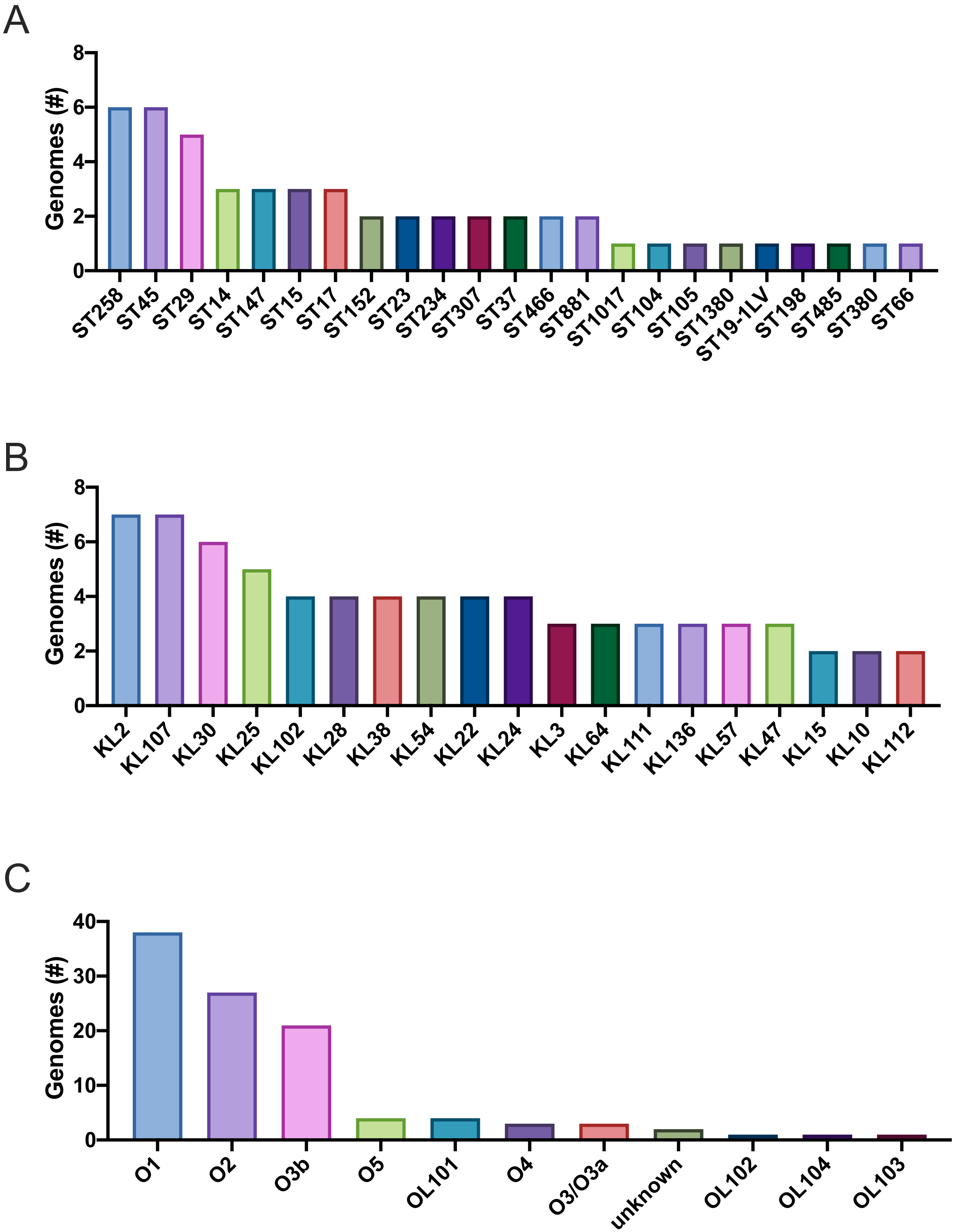

### Figure S3

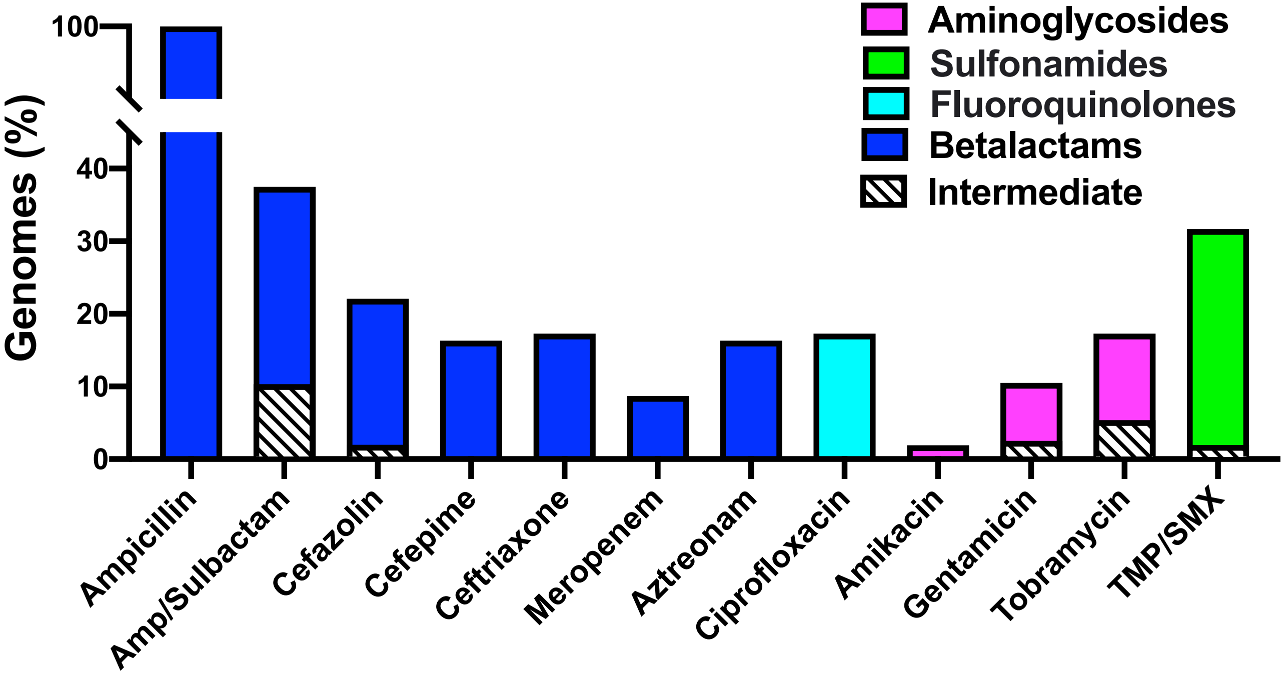

### Figure S4

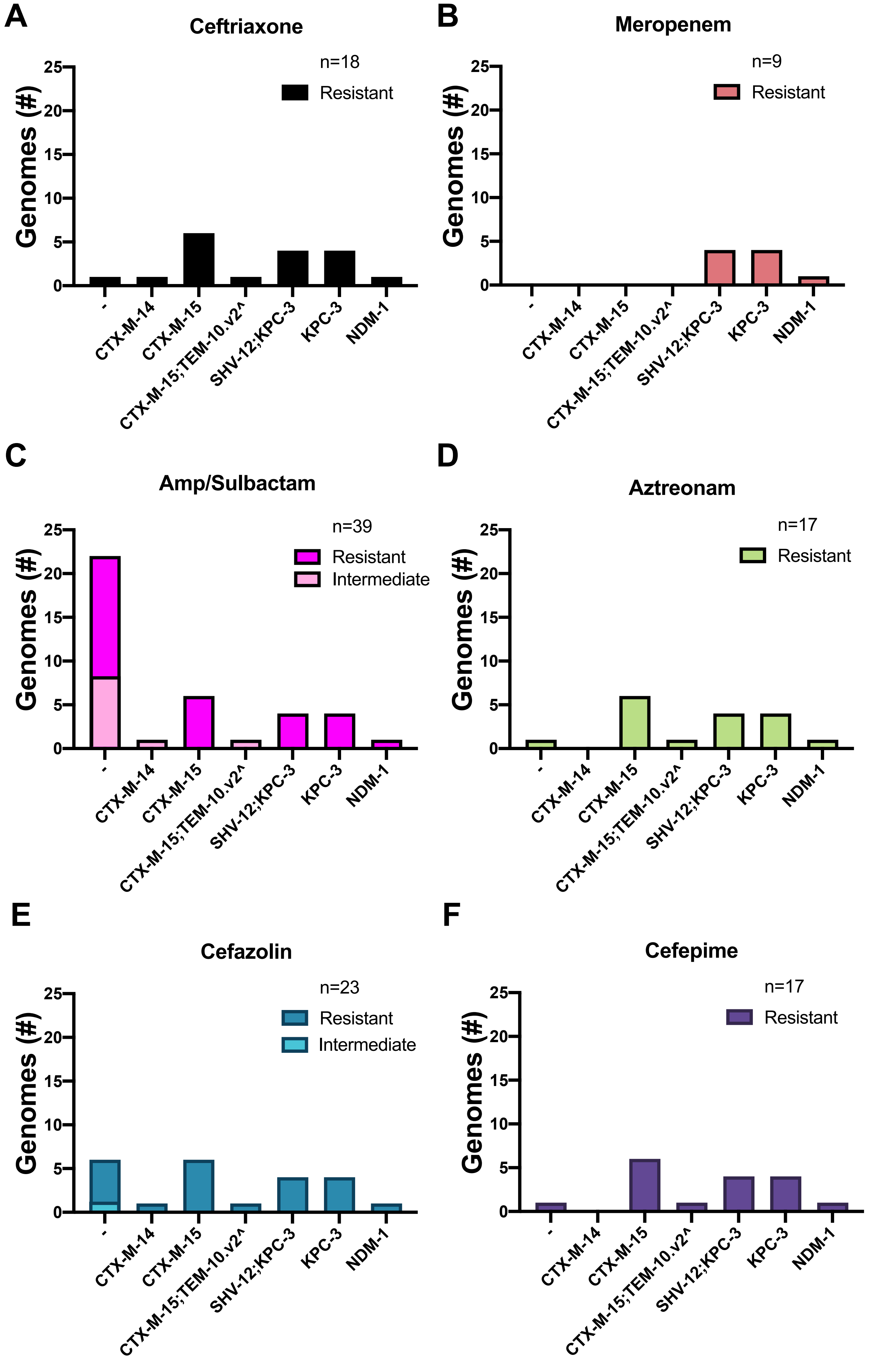

### Figure S5

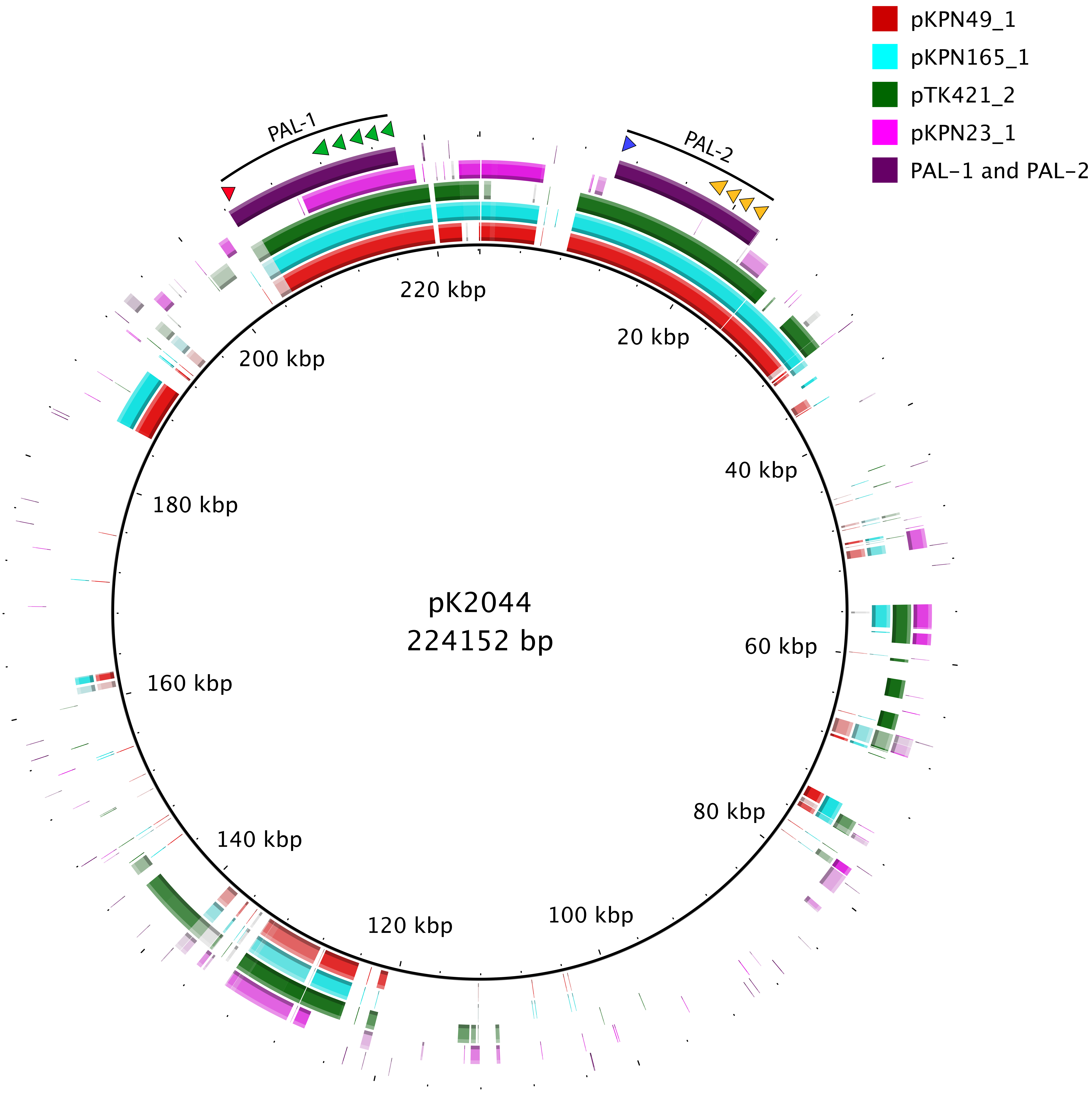

### Figure S6

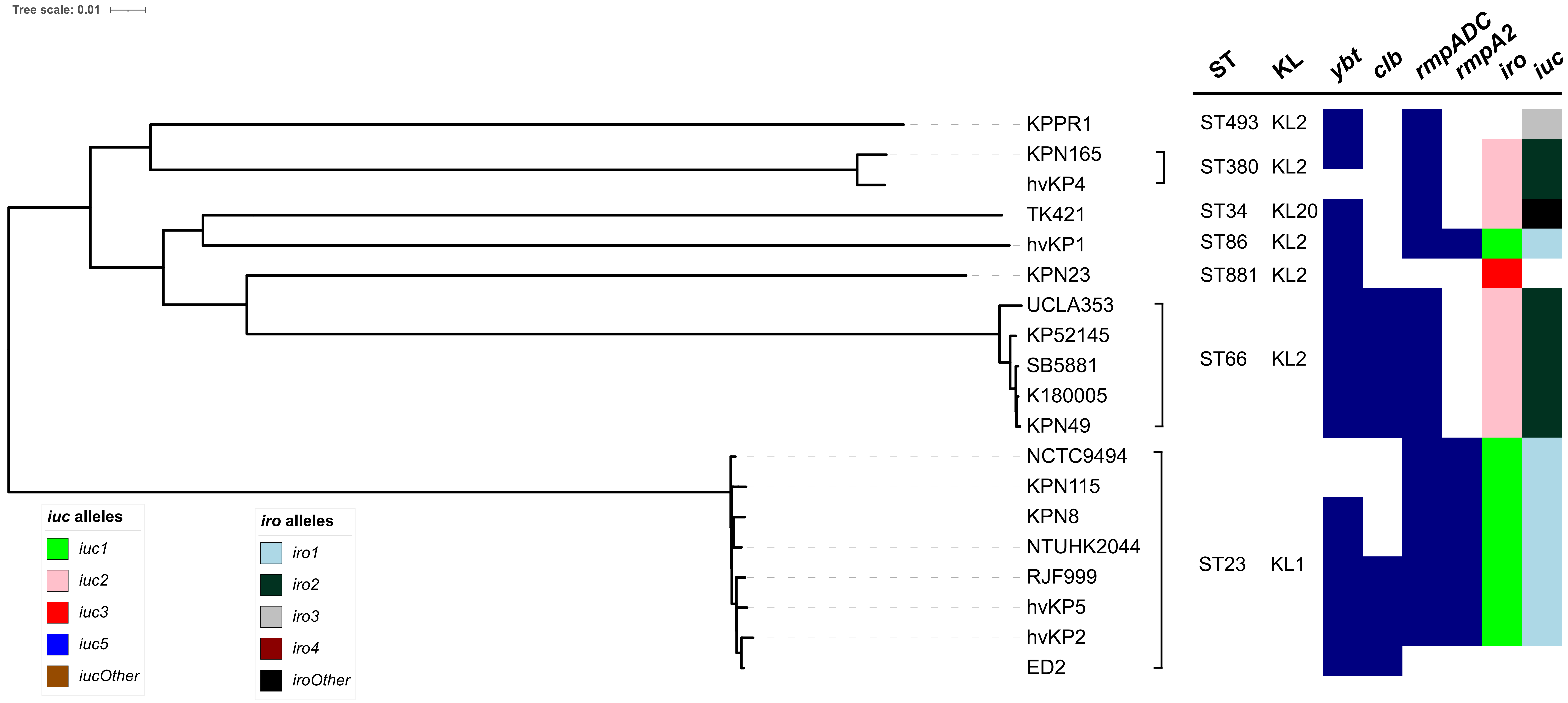
